# Structural basis for bacterial ribosome quality control

**DOI:** 10.1101/2020.09.03.281279

**Authors:** Caillan Crowe-McAuliffe, Hiraku Takada, Victoriia Murina, Christine Polte, Sergo Kasvandik, Tanel Tenson, Zoya Ignatova, Gemma C. Atkinson, Daniel N. Wilson, Vasili Hauryliuk

## Abstract

In all branches of life, stalled translation intermediates are recognized and processed by ribosome-associated quality-control (RQC) pathways. RQC begins with splitting of stalled ribosomes, leaving an unfinished polypeptide still attached to the large subunit. Ancient and conserved NEMF family RQC proteins target these incomplete proteins for degradation by the addition of C-terminal ‘tails.’ How such tailing can occur without the regular suite of translational components is, however, unclear. Using *ex vivo* single-particle cryo-EM, we show that C-terminal tailing in *Bacillus subtilis* is mediated by NEMF protein RqcH in concert with YabO, a protein homologous to, yet distinct from, Hsp15. Our structures reveal how these factors mediate tRNA movement across the ribosomal 50S subunit to synthesize polypeptides in the absence of mRNA or the small subunit.

## Introduction

In all cells, translational stalling on truncated or damaged mRNAs is harmful because it sequesters ribosomes from active protein production and can result in synthesis of cytotoxic truncated proteins. Therefore, ribosome-associated quality-control (RQC) pathways have evolved in all domains of life to disassemble such stalled complexes (Inada, 2020; Joazeiro, 2019). In eukaryotes, stalled 80S ribosomes are recognized and split into small 40S and large 60S subunits by Pelota/Dom34 and ABCE1/Rli1 (Franckenberg et al., 2012). The resulting 60S-peptidyl-tRNA complexes are then processed by the RQC pathway, where conserved NEMF-family proteins – Rqc2p in yeast and NEMF in humans – facilitate the addition of C-terminal alanine and threonine (CAT) tails to the nascent polypeptide chains (Brandman et al., 2012; Defenouillere and Fromont-Racine, 2017; Inada, 2020; Joazeiro, 2019; Kostova et al., 2017; Shen et al., 2015; Sitron and Brandman, 2019; Yan and Zaher, 2019). The nascent polypeptides are ubiquitinylated by Listerin/Ltn1 (Bengtson and Joazeiro, 2010; Lyumkis et al., 2014), and are extracted by p97/Cdc48 prior to proteasomal degradation (Defenouillere et al., 2013; Verma et al., 2013).

Bacterial NEMF-family homologs are members of the FbpA (fibronectin binding protein A) family of virulence factors. FbpA proteins from a number of Gram-positive bacterial species including *Enterococcus faecalis* (Singh et al., 2015), *Listeria monocytogenes* (Osanai et al., 2013), *Streptococcus pneumoniae* (Pracht et al., 2005) and *Bacillus subtilis* (Rodriguez Ayala et al., 2017) were proposed to directly mediate bacterial adhesion to the extracellular matrix, although a direct experimental demonstration of this function has been lacking. A recent study demonstrated that the *B. subtilis* NEMF homologue, RqcH (‘bacterial Rqc2 homolog’), is a *bona fide* bacterial RQC factor (Lytvynenko et al., 2019). *B. subtilis* RqcH is recruited to 50S-peptidyl-tRNA complexes to promote the addition of C-terminal polyalanine tails to stalled aberrant polypeptides, targeting incomplete proteins for degradation by the ClpXP machinery (Lytvynenko et al., 2019). Although *rqcH* is absent in a number of bacterial lineages, this discovery, along with the broad distribution of this factor in archaea, implies that the NEMF proteins were present in LUCA and that C-terminal tailing is therefore integral to RQC in all three domains of life (Burroughs and Aravind, 2014; Lytvynenko et al., 2019).

NEMF-family proteins are widely distributed in all three kingdoms of life and typically contain, from N-to C-terminus, (i) an NFACT-N domain followed by (ii) two helix-hairpin-helix (HhH) motifs, which have homology to DNA glycosylases but have no known enzymatic activity; (iii) a coiled-coil motif, consisting of two long α-helices separated by a small ‘middle’ domain, termed CC-M; (iv) an NFACT-R domain, which is predicted to bind RNA; and, (v) an NFACT-C domain of unknown function which is absent in bacterial NEMF homologs (Burroughs and Aravind, 2014; Shao et al., 2015). An early structural study proposed that yeast Rqc2p is bound to the 60S subunit around the P-site, likely recognising the peptidyl-tRNA (Lyumkis et al., 2014). Subsequently, two more structures of NEMF proteins bound to the large ribosomal subunit were reported: a yeast Rqc2p-60S complex with tRNAs bound in A- and P-sites (Shen et al., 2015), and an *in vitro* reconstituted mammalian 60S-Listerin-NEMF complex with a P-site peptidyl-tRNA (Shao et al., 2015). In both structures, the NFACT-N and HhH domains bound the stalled 60S complex close to the P-tRNA, and the coiled-coil spanned the A-site to contact the stalk base with the middle CC-M domain. Although these structures revealed the global binding mode of NEMF factors to the 60S ribosome and associated peptidyl-tRNA, due to the low resolution of the NEMF-family proteins at most only a partial molecular model could be built (Lyumkis et al., 2014; Shao et al., 2015; Shen et al., 2015). A detailed understanding of how NEMF-family proteins interact with RQC complexes, mechanistic insight into how NEMF homologues catalyze C-terminal tailing, and the identity of other factors involved in the bacterial RQC pathway have so far remained elusive.

Here we present *ex vivo* cryo-EM structures of *B. subtilis* RqcH bound to a 50S-peptidyl tRNA complex, and discover an additional factor, YabO, which was previously not known to be associated with RQC and which is co-distributed with RqcH across many bacterial lineages. Surprisingly, our series of RQC structures mimic distinct pre- and post-translocational states observed during canonical translation elongation. This provides the structural and mechanistic basis for how RqcH and YabO cooperate to mediate tRNA movement, and thereby processive alanine tailing, through an RQC translation cycle that is independent of mRNA, the small ribosomal subunit, and the translocase EF-G.

## Results

### Cryo-EM structures of *ex vivo* RqcH-50S complexes

To investigate how RqcH mediates C-terminal tailing in *B. subtilis*, we have determined cryo-EM structures of *ex vivo* RqcH-50S complexes. The RqcH-50S complexes were purified by affinity chromatography from *B. subtilis* cells expressing RqcH C-terminally tagged with a FLAG3 epitope (**Figure S1A**). Single particle cryo-EM analysis of the RqcH-50S complexes, with extensive *in silico* sorting, yielded four distinct 50S functional states, States A–D (**Figure S1B**). State A contained RqcH and a peptidyl-tRNA in an A/P-like configuration (**Figure 1A**), whereas State B contained RqcH, a peptidyl-tRNA in a classical P-site-like conformation, as well as an additional protein factor, which we identified as YabO (**Figure 1B, Figure S1B**) — a homolog of *E. coli* Hsp15. State C was similar to State A, but with the additional presence of a tRNA in the E-site (**Figure 1C**). State D contained YabO and P-site tRNA, but no RqcH, implying that RqcH had dissociated during sample preparation (**Figure S1B**). States A, B, C, and D were refined to average resolutions of 3.5 Å, 2.9 Å, 3.2 Å, and 2.6 Å, respectively (**Figure S2A-D**); however, while the 50S was well-resolved, the quality of the density for the ligands varied (**Figure S2E-H**). RqcH exhibited high flexibility in States A and C where the peptidyl-tRNA was in the A/P-state, but was better ordered in State B where YabO was present and the peptidyl-tRNA was in a classical P-site conformation (**Figure 1A-C** and **Figure S2E-G**). We further improved the cryo-EM map density for RqcH using multibody refinement (**Figure S1B**). The resulting map was sufficient for unambiguous fitting of individual domains of a homology model for *B. subtilis* RqcH based on the X-ray structures of RqcH homologs from related Gram-positive bacteria (Manne et al., 2019; Musyoki et al., 2016) (**Figure 1D, Table 1,** and **Video S1**). The NFACT-N and CC-M domains were relatively well resolved (**Figure S3A-C**), consistent with the presence of density for many bulky and aromatic sidechains, with the exception of the vestigial M domain, which was small and resembled a hairpin (**Figure S3D-G**). The NFACT-R and HhH domains appeared more flexible and less well-resolved (**Figure S3A-B**). Nonetheless, and with the exception of three short loops, we were able to model residues 2 to 565 of RqcH and use this model to fit and refine structures of RqcH in States A and B (**Table 1**).

**Figure 1.**
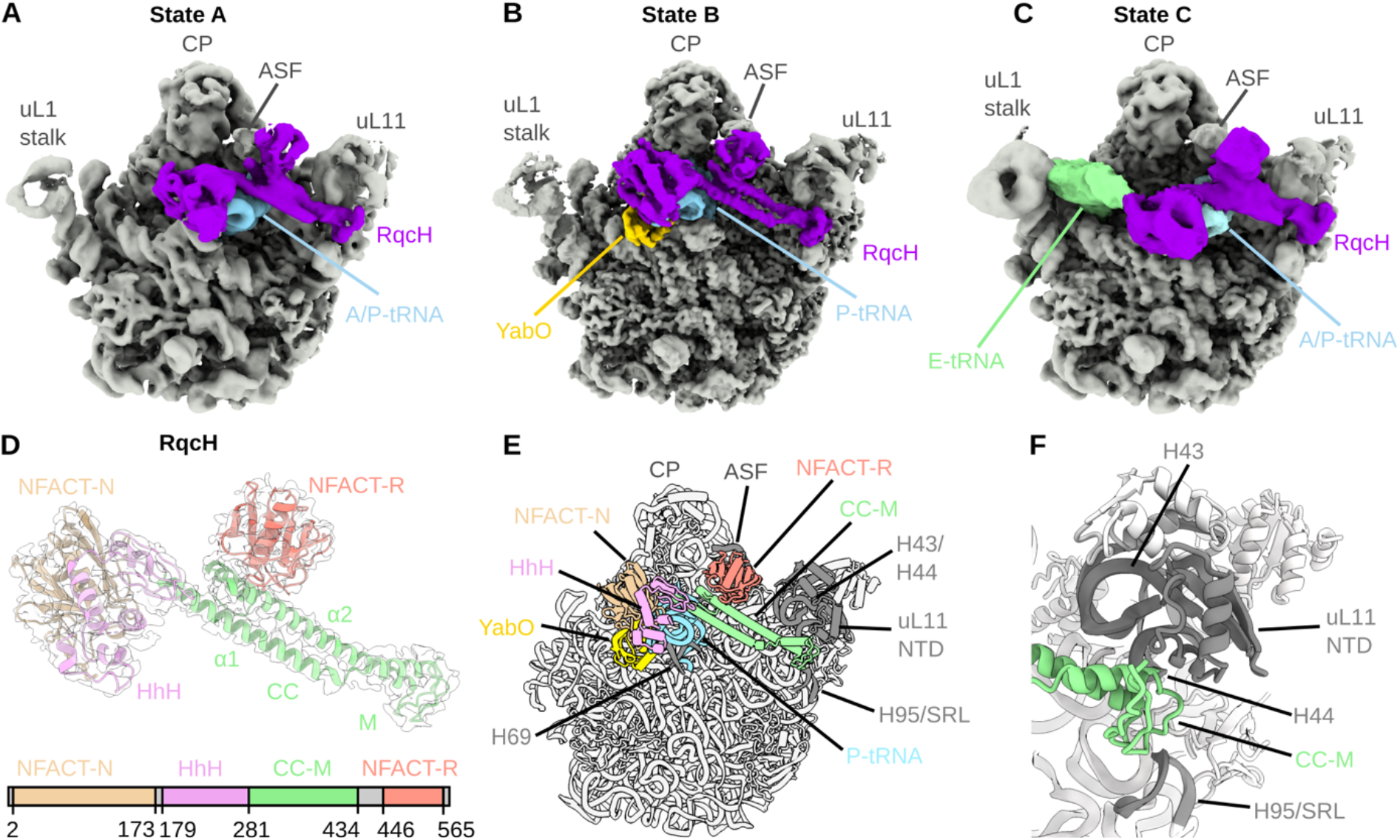
Cryo-EM structures of *B. subtilis* RqcH-50S complexes. (A-C) Cryo-EM maps of RqcH-50S complexes with (A) A/P-tRNA (State A), (B) P-tRNA, and YabO (State B), and (C) A/P-tRNA and E-tRNA (State C). 50S, grey; RqcH, purple; A/P- and P-tRNAs, light blue; YabO, yellow; and E-tRNA, green. (D) Cryo-EM map of RqcH from State B multibody refinement with RqcH model coloured by domain, according to the key (below). (E) Model of State B with RqcH domains labelled as in (D). (F) Highlight of the interaction between the CC-M domain of RqcH (green) and the uL11 stalk base and sarcin-ricin-loop (SRL).

**Table 1.** Cryo-EM data collection, model refinement and validation statistics.

| Collection details | RqcH collection 1 | RqcH collection 2 | YabO collection |
| --- | --- | --- | --- |
| Number of micrograph movies | 3 032 | 4 145 | 5 416 |
| Electron Fluence | 28.3 | 29.7 | 27.4 |
| Defocus range ($\mu\text{m}$) | −0.7–1.9 $\mu\text{m}$ | −0.7 to −1.9 | −0.4–1.9 $\mu\text{m}$ |
| Model composition | RqcH State A<br>(EMDB XXXX)<br>(PDB XXXX) | RqcH State B<br>(EMDB XXXX)<br>(PDB XXXX) | State B Multibody<br>refinement<br>(EMDB XXXX)<br>(PDB XXXX) |
| Average map resolution ($\text{\AA}$) | 3.5 | 2.9 | 4.0 |
| Number of particles | 10 703 | 74 210 | 74 210 |
| Non-hydrogen atoms | 93 138 | 93 796 | 8 157 |
| Protein residues | 3 720 | 3 809 | 751 |
| RNA bases | 2 999 | 2 997 | 118 |
| Refinement |  |  |  |
| Map CC around atoms | 0.82 | 0.90 | 0.73 |
| Map CC whole volume | 0.81 | 0.91 | 0.73 |
| Map sharpening B factor ($\text{\AA}^2$) | −66.07 | −71.99 | −98.32 |
| R.M.S. deviations |  |  |  |
| Bond lengths ($\text{\AA}$) | 0.009 | 0.017 | 0.008 |
| Bond angles ($^\circ$) | 1.136 | 1.151 | 1.010 |
| Validation |  |  |  |
| MolProbity score | 2.05 | 1.76 | 1.67 |
| Clash score | 7.78 | 4.45 | 4.22 |
| Poor rotamers (%) | 0.13 | 0.03 | 0.18 |
| Ramachandran plot |  |  |  |
| Favored (%) | 90.70 | 90.49 | 92.65 |
| Allowed (%) | 9.11 | 9.40 | 7.35 |
| Disallowed (%) | 0.17 | 0.11 | 0 |

### Interaction of RqcH on the 50S subunit

In the best-resolved state, State B, the RqcH N-terminal NFACT-N and HhH domains are located near the central protuberance (CP) between the P- and E-sites, while the coiled coils of the CC-M domain span the interface of the 50S subunit to the uL11 stalk base and then back to the A-site finger (ASF, H38), where the NFACT-R domain is positioned (**Figure 1E** and **Video S1**). The overall binding site of *B. subtilis* RqcH on the 50S subunit is similar to that of the eukaryotic homologues, yeast Rqc2p (Shen et al., 2015) and human NEMF (Shao et al., 2015), on the 60S subunit (**Figure S3H-O**), however, the higher resolution of RqcH-50S complexes reveals many additional details not observed before. The NFACT-N and HhH domains of RqcH interact predominantly with the anticodon-stem loop (ASL) of P-site tRNA (**Figure 1E** and **S3G**) and do not appear to establish any contact with the 50S subunit. This rationalizes how the previously reported D97/R98 (DR) and E121/I122/M123 (EIM) mutations in the NFACT-N domain specifically abrogate tRNA binding (Lytvynenko et al., 2019), since these motifs are located within loops that approach the anticodon of the tRNA (**Figure 2A-C**). In good agreement with a previous report (Lytvynenko et al., 2019), introduction of the DR and EIM mutations did not destabilise the interaction of RqcH with the 50S (**Figure 2E**). Indeed, one volume obtained from *in silico* sorting resembled State B (termed hereafter as State B*) but with little density for the NFACT-N and HhH domains, indicating that these domains can be flexible on the 50S and are not required for RqcH binding (**Figure S1B**). There are only two direct contacts between RqcH and components of the 50S subunit, namely between the RqcH NFACT-R domain and the ASF (**Figure 1E**) and between the distal portion of the RqcH CC-M domain and uL11/H44 at the stalk base (**Figure 1F**). Both interactions are necessary for RqcH function because (i) mutations in the conserved DWH motif of the NFACT-R domain, which is in close proximity to the ASF, leads to a loss in interaction with the 50S subunit as assessed by sucrose gradient centrifugation of cellular lysates (**Figure 2D-E**), and (ii) treatment of *B. subtilis* lysates with thiostrepton, an antibiotic that has an overlapping binding site with the CC-M domain (**Figure 2F-I**) abrogated the association of RqcH the 50S subunit (**Figure 2J**). In good agreement with a recent study of yeast Rqc2p (Osuna et al., 2017), antibiotics targeting the peptidyl-transferase center (lincomycin), the small ribosomal subunit (viomycin), or canonical GTPase translation factors EF-Tu (kirromycin) and EF-G (fusidic acid) did not perturb RqcH association with the 50S (**Figure 2J**). Puromycin, which releases the nascent chain if the A-site is accessible, had only a mild effect, in any (**Figure 2J**).

**Figure 2.**
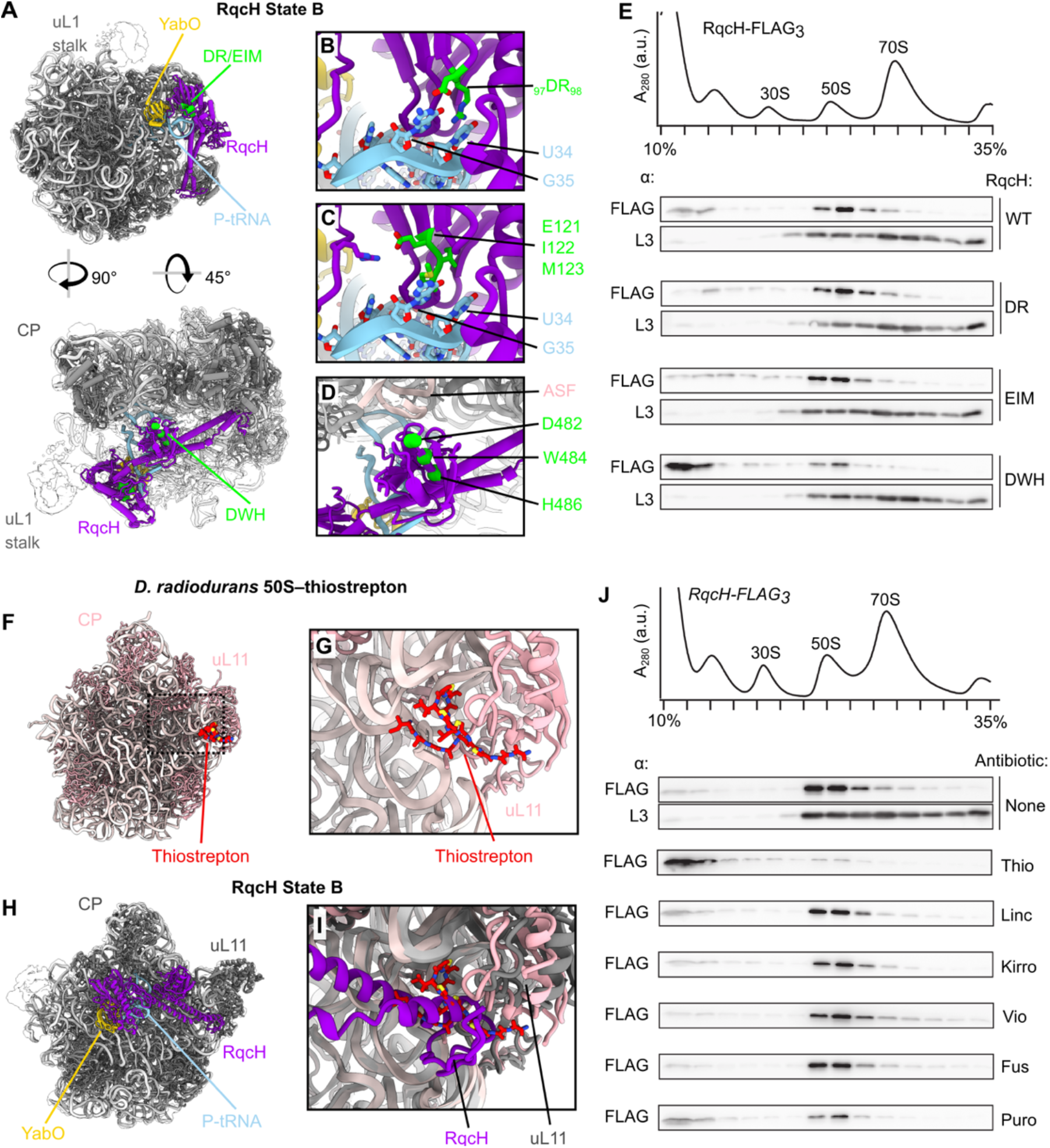
The interaction of RqcH with the 50S probed with mutagenesis and antibiotics. (A-D) Structural analysis of RqcH mutants; (B) DR, D97A/R98A, (C) EIM, E121A/I122A/M123A, and (D) DWH, D482A/W484A/H486A. Mutated residues are shown in green. (E) Sucrose gradient sedimentation of RqcH-FLAG3 mutants. Immunoblots were probed with either α-FLAG to detect RqcH wild-type and mutant RqcH variants, or α-L3. (F) The thiostrepton-bound 50S from *D. radiodurans* (PDB 3CF5) (Harms et al., 2008). (G) Close view of thiostrepton interacting with the ribosome close to uL11. (H) RqcH State B shown from the same perspective as (F). (I) Overlay of thiostrepton-bound 50S and RqcH state B, from the same view as (G). (J) Sucrose gradient sedimentation of RqcH in the presence or absence of translation-targeting antibiotics added after cell lysis: Thio, thiostrepton (50 μM); Kirro, kirromycin (50 μM); Linc, lincomycin, (1 μM); Vio, viomycin (100 μM); Fus, fusidic acid (100 μM); Puro, puromycin (1 mM).

### Discovery of a role for YabO during bacterial RQC

State B contained additional density, positioned between the RqcH NFACT-N domain, P-tRNA, and 23S rRNA helices 68 and 69, that did not correspond to RqcH or any ribosomal component (**Figure 3A, B**). This density was assigned to YabO based on mass spectrometry (**Table S1**) and the excellent agreement between the density features and a fitted homology model for *B. subtilis* YabO using the crystal structure of *E. coli* Hsp15 as the template (Staker et al., 2000) (**Figure 3C**). *E. coli* Hsp15 binds 50S-peptidyl-tRNA complexes (Korber et al., 2000) and can translocate the peptidyl-tRNA from the A-to the P-site (Jiang et al., 2009). While *E. coli* Hsp15 was reported to bind at the central protuberance of the 50S (Jiang et al., 2009), YabO instead binds the 50S at a distinct site adjacent to H69 (**Figure S3P-R**). *E. coli* and other gamma-proteobacteria do not contain RqcH homologs, suggesting that *E. coli* Hsp15 may function differently than *B. subtilis* YabO. Additionally, *E. coli* Hsp15 has a C-terminal extension (CTE) that is absent in YabO (**Figure 3D** and **S3S**). YabO/Hsp15 homologues across diverse bacterial clades divide into those either having the CTE, such as *E. coli* Hsp15, or not, such as YabO (**Figure 3D**), and strikingly this division is strongly associated with the presence or absence of RqcH. Presence of the Hsp15 CTE is entirely mutually exclusive with the presence of RqcH, and – with few exceptions – bacteria with YabO/Hsp15 homologues lacking the CTE contain RqcH (**Figure 3D** and **Table S2**). Together with the presence of Proteobacteria in both clades of YabO/Hsp15, this suggests these proteins are not functionally equivalent orthologues but are rather functionally divergent paralogues. This is further supported by the observation that, unlike Hsp15, expression of YabO is not induced by heat shock (Nicolas et al., 2012). Collectively, these findings suggest that YabO homologues are likely to be involved in RQC in bacteria containing RqcH, but raises the question as to the role of Hsp15 and its CTE in bacteria lacking RqcH.

**Figure 3.**
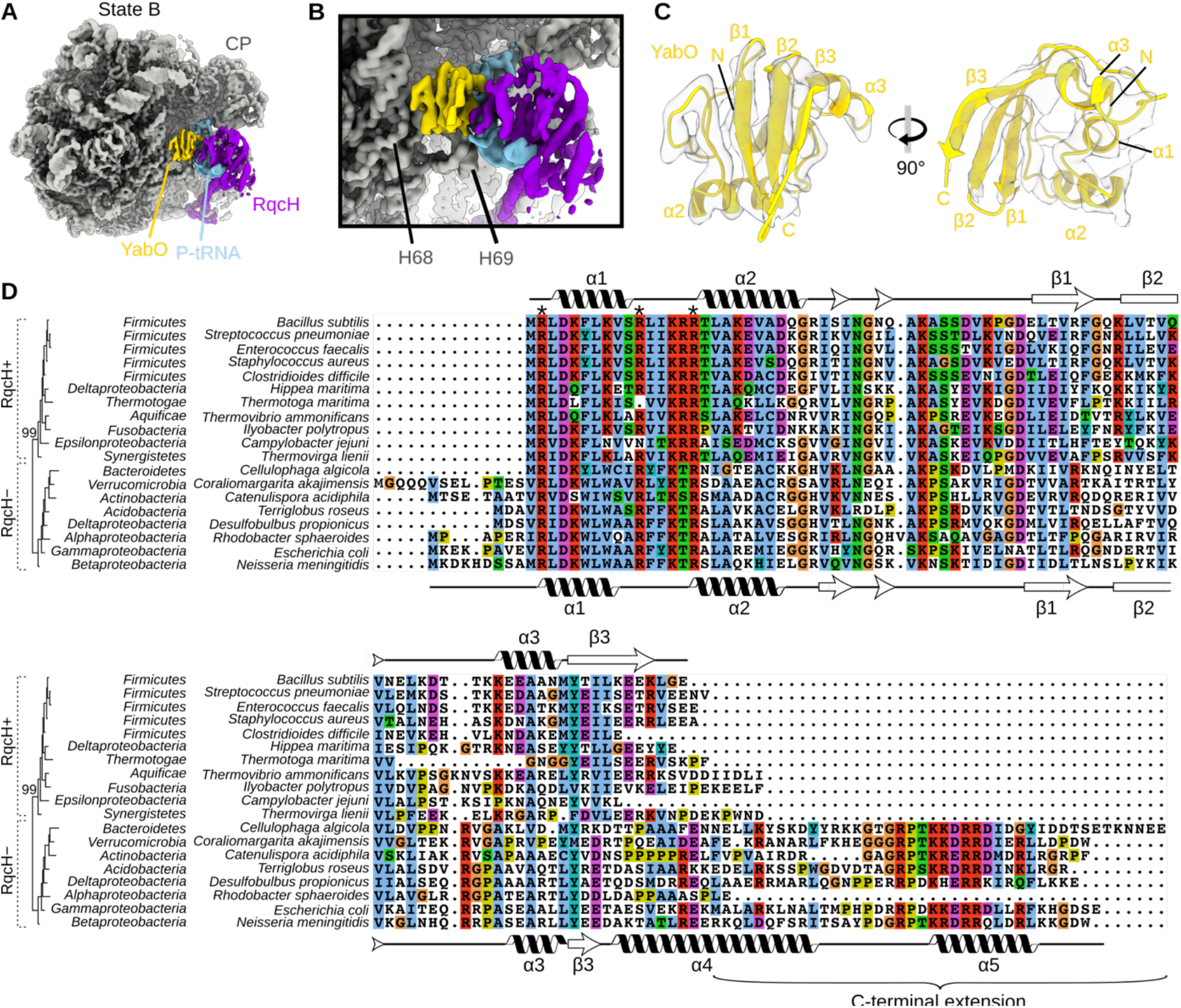
YabO binds the 50S adjacent to RqcH, and lack of the C-terminal extension in YabO/Hsp15 is associated with the presence of RqcH. (A-B) View of YabO in RqcH State B showing global position on the 50S (A) or zoomed view (B). (C) Views of YabO model fitted into density from state B. (D) Phylogenetic tree and sequence alignment of YabO/Hsp15 homologues from diverse bacteria. The RqcH+/RqcH-phylogenetic split is supported with 99% bootstrap support. Secondary structure, calculated with DSSP or predicted by PSI-PRED where no structure was available, is shown above (YabO, present study) or below (*E. coli* Hsp15, from PDB 1DM9 (Jiang et al., 2009)).

To validate the role of *B. subtilis* YabO in RQC we affinity-purified YabO with a C-terminal FLAG3-tag, yielding YabO-50S complexes (**Figure S4A**). RqcH co-purified with these complexes, as confirmed by mass spectrometry (**Table S1**). Single particle cryo-EM analysis and *in silico* sorting yielded State B with RqcH and P-site tRNA as well as State D with P-site tRNA but no RqcH (**Figure S4B**), both of which were also observed for the RqcH pull-outs (**Figure 1B** and **S1B**). We observed an additional novel State E, containing YabO with P- and E-site tRNAs, but no RqcH (**Figure S4B**). States B, D and E in the YabO pull-out dataset were refined to average resolutions of 3.2 Å, 2.6 Å and 3.2 Å, respectively (**Figure S5A-C**). In most states YabO was well resolved (**Figure S5D-F**) and established defined contacts with H69 of the 50S and the ASL of the P-site tRNA (**Figure 4A-D**). The interaction between YabO and RqcH was less well-resolved, and does not appear to be essential for recruitment of these factors to the 50S since RqcH migrated with the 50S in the absence of YabO and *vice versa* (**Figure 4E**). By contrast, interaction with 23S rRNA H69 is critical for YabO function, since mutation of the conserved Arg16 to Ala (R16A) completely abolished YabO association with the 50S subunit (**Figure 3D, 4E, S4A,** and **Table S1**). Comparison of State A (RqcH and A/P-site tRNA, **Figure 4F**) with State B (RqcH with YabO and a P-tRNA, instead of an A/P-tRNA, **Figure 4G**) suggests that YabO participates in the translocation of the peptidyl-tRNA from the A-site (**Figure 4H**) into the P-site (**Figure 4I**), consistent with previous proposals (Jiang et al., 2009). The entire RqcH protein shifted with the tRNA during the translocation event, with large-scale movements in the range of 15–20 Å (**Figure 4J** and **Video S2**). The shift in RqcH is also accompanied by a corresponding movement in the uL11 stalk base to which RqcH is tethered via the CC-M domain (**Figure 4J**).

**Figure 4.**
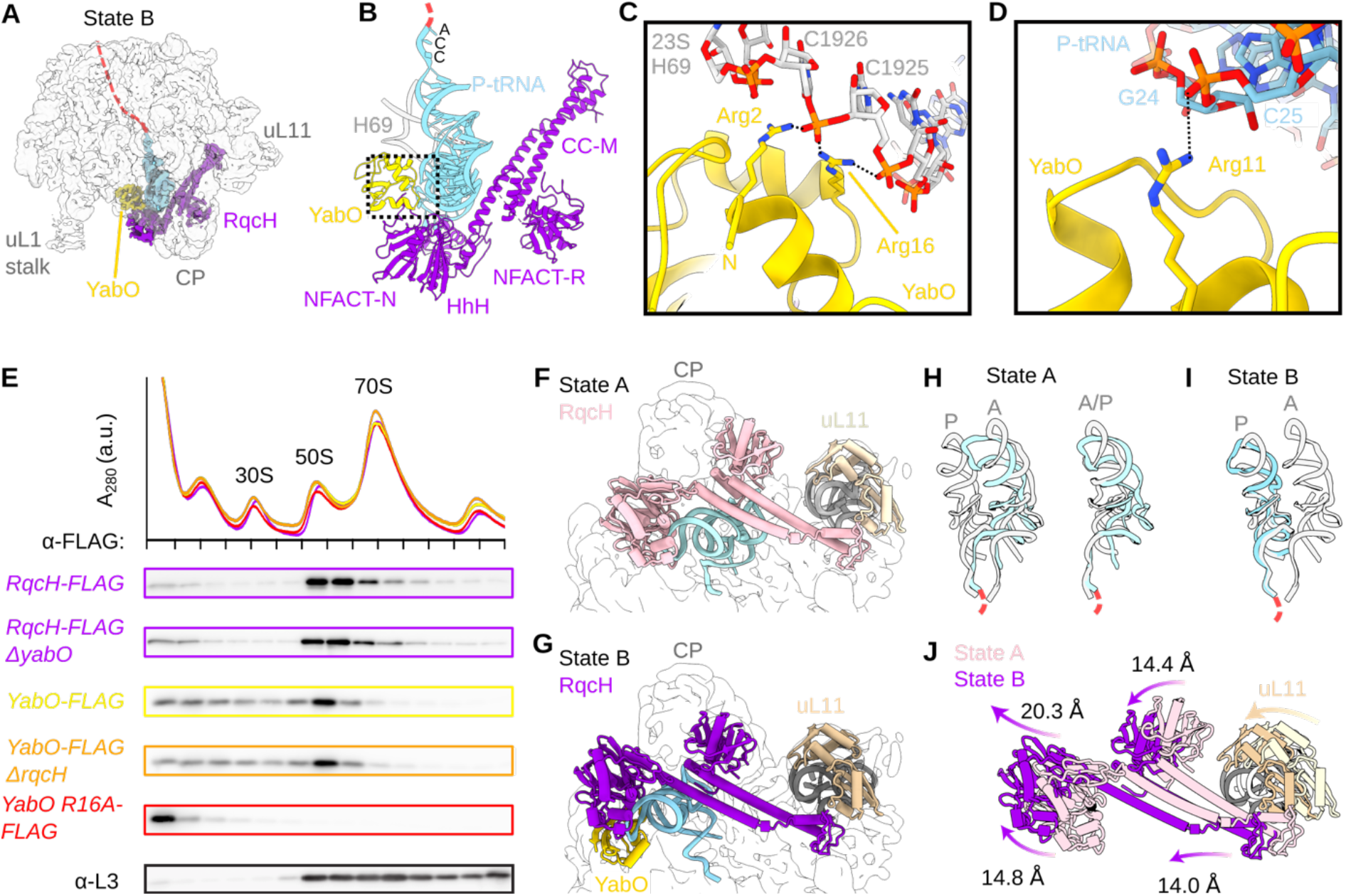
YabO stabilizes a classical P-site tRNA conformation. (A and B) Overview of YabO (yellow) interactions within State B. (C and D) Selected interactions between YabO and 23S H69 (grey, C) or the P-site tRNA (light blue, D). (E) Sucrose gradient analysis of *B. subtilis* strains expressing FLAG-tagged RqcH and YabO. Fractions were analysed by immunoblot with α-FLAG or α-L3. (F and G) Comparison of RqcH and tRNA within (F) State A and (G) State B. (KH and I) Comparison of tRNAs (cyan) from (H) State A or (I) State B with classical A- and P-site tRNAs (PDB 6CFJ) (Tereshchenkov et al., 2018) or hybrid A/P-site tRNA (PDB 6R6P) (Shanmuganathan et al., 2019). (J) Superposition of RqcH from states A (pink) and B (purple) with degrees of movement indicated. uL11 and the stalk base is shown for reference.

### Identification of RsfS in complex with RqcH-50S complexes

During 3D classification of the 50S subunits we also noticed a substoichiometric extra density in the vicinity of uL14. This density was further improved by focussed classification (**Figure S4B**) and identified as *B. subtilis* ribosomal silencing factor RsfS, bound in a position analogous to that observed previously on the 50S subunit (Brown et al., 2017; Khusainov et al., 2020; Li et al., 2015) (**Figure S6A-E**). This assignment is supported by mass spectrometry (**Table S1**) and retrospective inspection of the RqcH-pullout also revealed substoichiometric density for RsfS on the 50S subunit (**Figure S6C**). RsfS prevents association of 50S and 30S subunits (Hauser et al., 2012; Khusainov et al., 2020; Li et al., 2015) and therefore its presence in our datasets may indicate that RsfS also plays a similar role during RQC, analogous to that of Tif6/eIF6 in eukaryotic RQC (Su et al., 2019) (**Figure S6F**). Curiously, in the RqcH pull-out dataset we observed formation of 50S disomes (**Figure S1B**) containing RqcH, YabO and P-tRNA, i.e. dimerization of State B (**Figure S6G,H**). While an overlay of the 50S-bound RsfS with the structure of the 50S disome reveals that RsfS would prevent 50S dimerization (**Figure S6I-J**), it remains to be determined whether this has any physiological relevance.

### Interaction of RqcH with tRNA^Ala^ on the ribosome

Although the P-site tRNA in State B was relatively well-resolved, the resolution was not sufficient to unambiguously distinguish the tRNA species. To test whether the sample contained *bone fide* RQC complexes containing tRNA^Ala^ we performed tRNA microarray analysis on the RqcH pull-out sample. As expected, we observed an enrichment for tRNA^Ala(UGC)^ in our samples over the lysate. However, we also observed an enrichment of tRNA^Ala(IGC)^ (**Figure 5A**), which was not detected previously in the bacterial RqcH-complexes (Lytvynenko et al., 2019), but was observed in eukaryotic Rqc2p-60S complexes (Shen et al., 2015). Our findings suggest that *B. subtilis* RqcH can selectively recruit both Ala-tRNA^Ala^ isoacceptors to the ribosome to synthesize alanine tails. In State B, although the tRNA^Ala^ is bound to the 50S similarly to a P-site tRNA, the ASL element undergoes dramatic rearrangements (**Figure 5B,C**). Specifically, the ASL is unwound compared to a classical ASL-helix conformation and the anticodon nucleotides 34-36 are splayed apart and poorly ordered (**Figure 5C**). Arg125 from the NFACT-N domain of RqcH inserts into the ASL, where it interacts with the nucleotide in position 32, which is U32 in tRNA^Ala(UGC)^ and C32 in tRNA^Ala(IGC)^ (**Figure 5D**). We doubt that the Arg125 interaction contributes to defining tRNA specificity of RqcH since many *B. subtilis* tRNAs have either U or C at position 32. No state contained both A- and P-site tRNAs, indicating that, like in regular translation, peptide bond formation is fast, and following peptidyl transfer the tRNAs move rapidly into A/P- and E-sites. Comparison of State B (RqcH, YabO and P-site tRNA, **Figure 5E**) with State C (RqcH, A/P-tRNA and E-site tRNA, **Figure 5F**), suggests that YabO needs to dissociate from the 50S subunit to allow the uncharged P-site tRNA to move into the E-site (**Figure 5G** and **Video S3**). Similarly, RqcH would also need to rearrange to accommodate uncharged tRNA at the E-site, which involves a scissor-like separation of the coiled coils within the CC-M domain, such that the NFACT-N and HhH domains shift by an impressive 30 Å out of the E-site (**Figure 5H** and **Video S3**).

**Figure 5.**
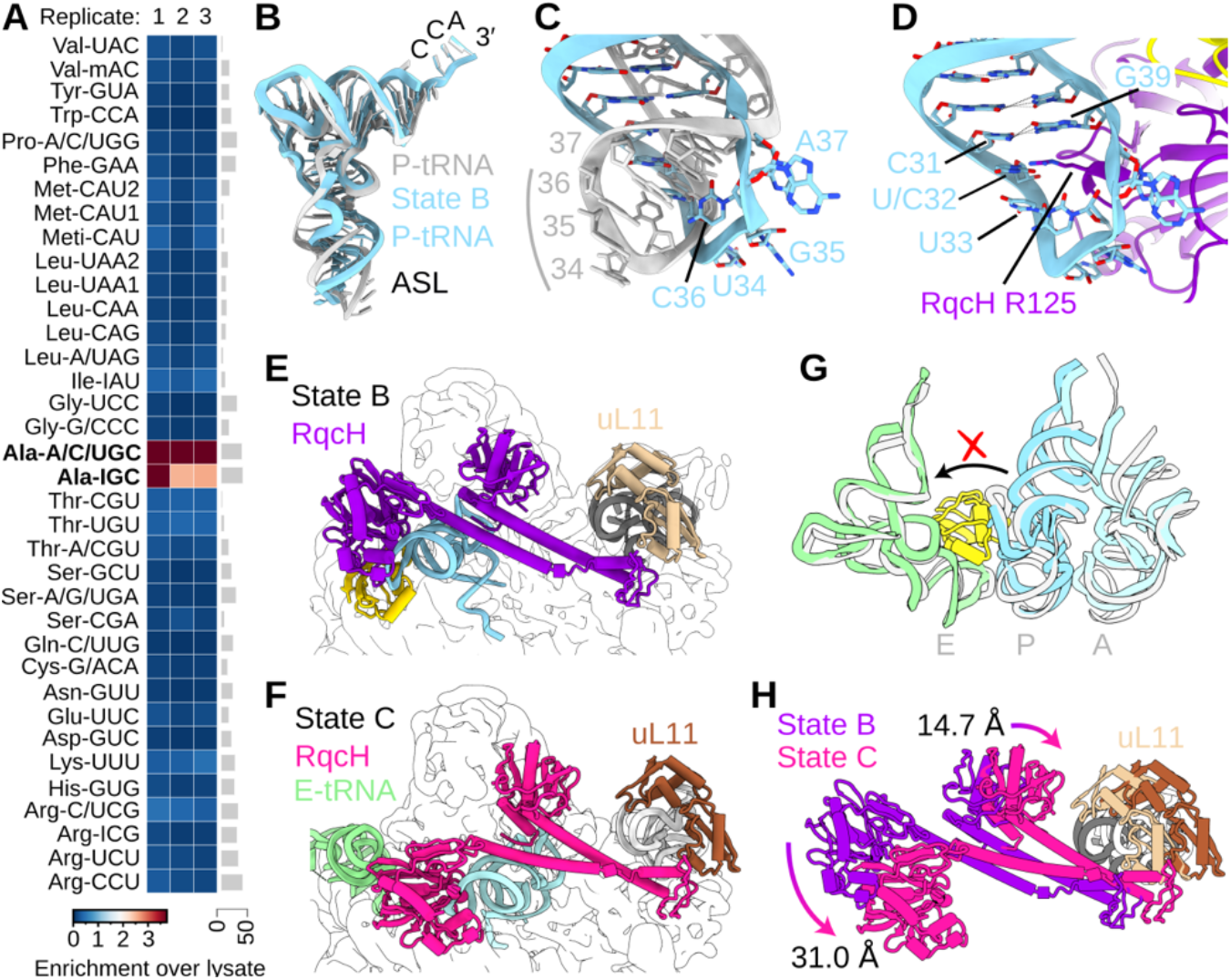
tRNA composition and dynamics within RqcH-50S complexes. (A) tRNA microarray analysis of the RqcH-50S sample. Three replicates are shown with the grey bars on the right representing an example of covariance analysis between replicate 1 and 3. Confidence intervals between replicate 1 and 2, 2 and 3 and 1 and 3 were 94%, 93% and 90%, respectively. Colour key indicates the fold-enrichment over lysate. (B-C) Comparison of RqcH-bound P-site tRNA from State B (blue) with canonical P-site tRNA from a 70S complex (grey, PDB 6CFJ) (Tereshchenkov et al., 2018). ASL, anticodon-stem loop. (D) Same view as (C) of the State B P-tRNA, but with RqcH (purple) and YabO (yellow) included. (E-F) Comparison of RqcH, tRNA and YabO within (E) State B and (F) State C. (G) Comparison of A/P-(cyan), P-(blue) and E-site (green) tRNAs from States B and C with canonical A-, P-, and E-site tRNAs (grey, (PDB 6CFJ) (Tereshchenkov et al., 2018). (H) Conformational changes of RqcH and the uL11 stalk base between States B and C.

## Discussion

Collectively, our ensemble of structures enables us to present a model for how polyalanine tailing of aborted 50S-peptidyl-tRNA complexes is catalysed by RqcH and YabO (**Figure 6** and **Video S4**). We suggest that dissociated 50S subunits with a peptidyl-tRNA are prevented from reassociation with the 30S by RsfS (**Figure 6A**). These 50S-peptidyl-tRNA complexes are recognized by YabO, which binds and stabilizes the peptidyl-tRNA in the P-site (State D, **Figure 6B**). This frees the A-site so that RqcH can deliver Ala-tRNA^Ala^. Following peptide-bond formation this results in a complex with uncharged tRNA at the P-site and the peptidyl-tRNA at the A-site (**Figure 6C**). To allow the uncharged tRNA to relocate to the E-site, YabO must dissociate from the 50S, thus permitting the peptidyl-tRNA to adopt an A/P-like configuration (State C, **Figure 6D**). Dissociation of the uncharged tRNA from the E-site of this complex then leaves a state with RqcH and A/P-tRNA (State A, **Figure 6E**) which allows rebinding of YabO, thus shifting the A/P-tRNA into the P-site (State B, **Figure 6F**) and completing the translocation cycle. Successive binding-dissociation cycles of YabO could act as a pawl of the RQC elongation ratchet, thus driving the processivity of alanine-tailing. Our observation of State E with YabO and P- and E-site tRNAs – but no RqcH – suggests that if YabO rebinds before E-tRNA release, then RqcH would dissociate, thus providing an alternative pathway back to State D (**Figure S5G**). State B*, which contains a partially dissociated RqcH (**Figure 4G**), indicates that RqcH may processively recruit new Ala-tRNAs while still tethered to the ribosome. Alternatively, RqcH could completely dissociate, leading back to State D (**Figure 6B**) and thereby requiring Ala-tRNA^Ala^ to be delivered to the A-site by another RqcH molecule.

**Figure 6.**
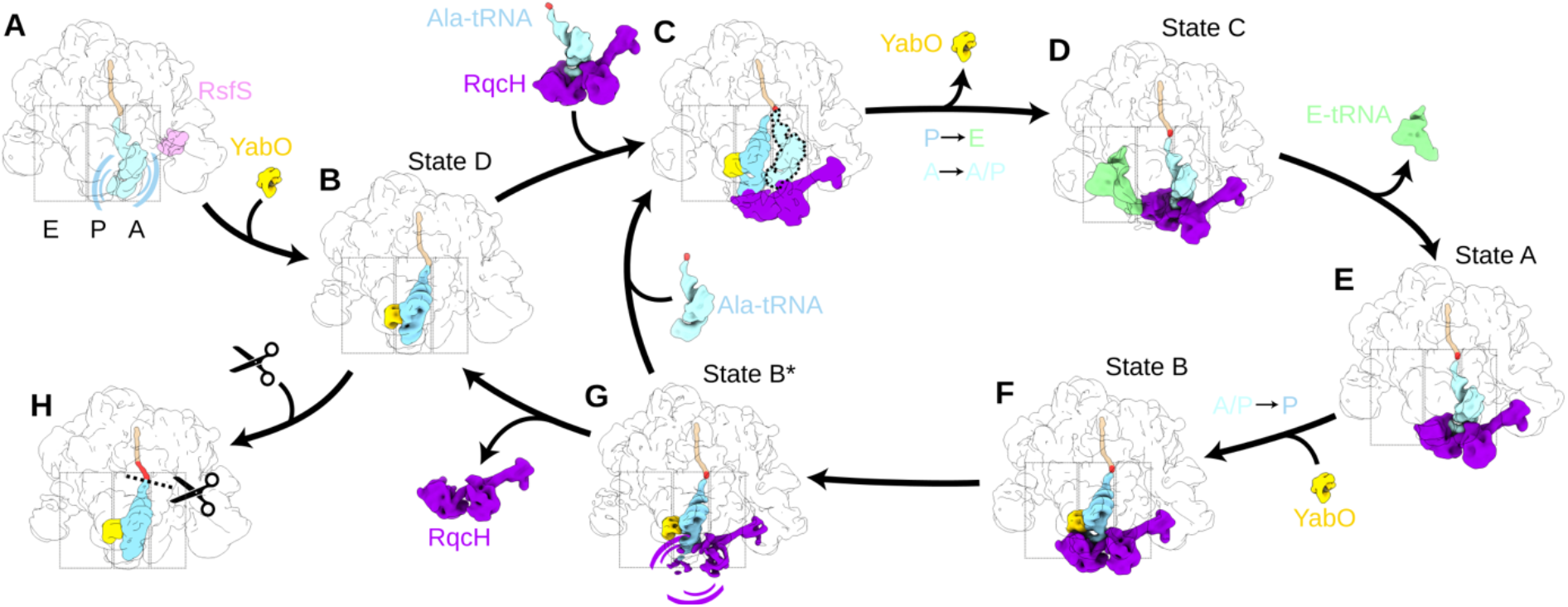
Model of C-terminal alanine tailing on the 50S mediated by RqcH and YabO. (A) An RQC substrate with 50S and flexible peptidyl-tRNA (with nascent chain shown in tan) is prevented from reassociation with the 30S by RsfS. (B) Binding of YabO to this 50S complex stabilizes the tRNA in the P-site (State D). (C) Hypothesized transient state in which Ala-tRNA^Ala^ is delivered to State D by RqcH. tRNA accommodates to the A-site followed by rapid peptidyl transfer. (D) YabO dissociation facilitates translocation-type movement of the tRNAs from P-to E- and A- to A/P-states (State C). (E) Dissociation of E-tRNA, results in a modest shift in the RqcH NFACT-N position (State A). (F) Binding of YabO stabilizes the tRNA in the classical P-site conformation, with concomitant movement of RqcH on the 50S (State B). (G) Partial dissociation of RqcH, evidenced by classes in which only the RqcH CC-M domain was observed (State B*). This can lead to either full RqcH dissociation (State D) or delivery of the next Ala-tRNA^Ala^, leading back to (C). (H) Hypothesized termination state, in which an unknown factor releases the alanine-tailed nascent polypeptide chain.

While our study provides structural insight into the mechanics of the bacterial RQC elongation cycle, a number of questions remain. Which cellular stresses and/or translational states lead to splitting of translating 70S ribosomes, and are there unknown factors that mediate this process? What is the functional state of the 50S-peptidyl-tRNA complex following splitting, and does it differ depending on the triggering conditions? How is polyalanine tail length regulated and eventually terminated? Is there a dedicated termination factor that mediates release of the tagged nascent polypeptide chain in bacteria, analogous to Vms1/ANKZF1 in eukaryotes? (Kuroha et al., 2018; Su et al., 2019; Verma et al., 2018; Yip et al., 2019; Zurita Rendon et al., 2018). In summary, we demonstrate the involvement of YabO in RqcH-mediated bacterial RQC, and propose an alternative model for protein synthesis on the ribosome that utilizes binding and rebinding of two non-GTPase protein factors to execute a whole elongation cycle without the small ribosomal subunit or mRNA.

## MATERIALS AND METHODS

### Bacterial strains

*B. subtilis* and *E. coli* strains (Crowe-McAuliffe et al., 2018; Guerout-Fleury et al., 1996; Guerout-Fleury et al., 1995; Horinouchi and Weisblum, 1982; Murina et al., 2019; Takada et al., 2014) used in this study are listed in **Table S3**. All *B. subtilis* strains used were derivatives of the wild-type 168 strain.

### DNA and plasmids

Plasmids as well as DNA oligonucleotides used in this study are listed in **Table S3**. All *B. subtilis* strains used were derivatives of the wild-type 168 strain. Mutant strains were constructed by transformation with plasmids or DNA fragments (the latter were generated by recombinant PCR, combining 3 or 4 smaller PCR fragments) and relied upon *in vivo* recombination, followed by selection for antibiotic resistance. The plasmids carried engineered *B. subtilis* genes of interest flanked by sequences corresponding to the integration target loci. PCR fragments and plasmids used in the study, as well as the schematics of their generation, are provided in **Table S3**. Plasmids and PCR fragments were constructed by standard cloning methods including PCR, Phusion Site-Directed Mutagenesis (Thermo Fisher Scientific) and Gibson assembly (NEB).

### Sucrose gradient fractionation and Western blotting

Sucrose gradient fractionation and Western blotting were carried out as described earlier (Takada et al., 2020), with minor modifications. *B. subtilis* strains were pre-grown on LB plates overnight at 30 °C. Fresh individual colonies were used to inoculate 200 mL LB cultures. The cultures were grown until OD600 of 0.8 at 37 °C and the cells were collected by centrifugation at 8,000 rpm for 5 minutes in JLA-16.25 rotor (Beckman Coulter), dissolved in 0.5 mL of HEPES:Polymix buffer, 5 mM Mg(OAc)2 (Takada et al., 2020) supplemented with 2 mM PMSF. Cells were lysed using FastPrep homogenizer (MP Biomedicals) by four 20 second pulses at speed 6.0 mp/sec with chilling on ice for 3 minutes between the cycles) and the resultant lysates were clarified by ultracentrifugation at 14,800 rpm for 20 minutes in F241.5P rotor using Microfuge 22R centrifuge (Beckman Coulter). 10 A260 units of each extract were loaded onto 10-35% (w/v) sucrose density gradients in Polymix buffer, 5 mM Mg(OAc)2. Gradients were resolved at 36,000 rpm for 3 hours at 4 °C in SW41 rotor (Beckman). Both separation and fractionation of gradients used a Biocomp Gradient Station (BioComp Instruments) with A280 as a readout.

For Western blotting, 0.5 mL fractions were supplemented with 1.5 mL of 99.5% ethanol, and precipitated overnight at −20 °C. After centrifugation at 14,800 rpm for 30 minutes at 4 °C the supernatants were discarded and the samples were dried. The pellets were resuspended in 40 μL of 2x SDS loading buffer (100 mM Tris-HCl pH 6.8, 4% SDS (w/v) 0.02% Bromophenol blue, 20% glycerol (w/v) 4% β-mercaptoethanol), resolved on the 12% SDS PAGE and transferred to nitrocellulose membrane (Trans-Blot Turbo Midi Nitrocellulose Transfer Pack, Bio-Rad, 0.2 μm pore size) using Trans-Blot Turbo Transfer Starter System (Bio-Rad) (10 minutes, 2.5A, 25V). Membranes were blocked for one hour in PBS-T (1 × PBS, 0.05% Tween-20) with 5% w/v non-fat dry milk at room temperature. RqcHFLAG3 was detected using anti-Flag M2 primary (Sigma-Aldrich, F1804; 1:10,000 dilution) antibodies combined with anti-mouse-HRP secondary (Rockland; 610-103-040; 1:10,000 dilution) antibodies. Ribosomal protein L3 was detected using anti-L3 primary antibodies (a gift from Fujio Kawamura; 1:20,000 dilution) combined with goat anti-rabbit IgG-HRP secondary antibodies (Sigma-Aldrich, A0545; 1:10,000 dilution). ECL detection was performed using WesternBright^™^ Quantum (K-12042-D10, Advansta) Western blotting substrate and ImageQuant LAS 4000 (GE Healthcare) imaging system.

### Immunoprecipitation of FLAG3-tagged proteins

Strains expressing FLAG3-tagged proteins were pre-grown on LB plates overnight at 30 °C. Fresh individual colonies were used for inoculation and grown in LB medium. 3× 1 L cultures were grown at 37 °C to OD600 = 0.8. Cells were collected by centrifugation (8 000 rpm for 10 min at 4 °C, JLA-16.25 Beckman Coulter rotor), pellets frozen in liquid nitrogen and stored at −80°C. Cell pellets were resuspended in 8 mL of cell opening buffer (95 mM KCl, 5 mM NH4Cl, 20 mM Hepes (pH = 7.5), 1 mM DTT, 15 mM Mg(OAc)2, 0.5 mM CaCl2, 8 mM putrescine, 1 mM spermidine, 1 tablet of cOmplete™ EDTA-free Protease Inhibitor Cocktail (Roche) per 50 mL of buffer) and disrupted using FastPrep homogeniser (MP Biomedicals) with 0.1 mm Zirconium beads (Techtum) in 6 cycles by 20 seconds with 3 minute chill on ice. Cell debris was removed by centrifugation at 14,800 rpm for 20 minutes 4 °C in F241.5P rotor using 149 Microfuge 22R centrifuge (Beckman Coulter). The supernatant was combined with 100 μL of ANTI-FLAG M2 Affinity Gel (Sigma) pre-equilibrated in cell opening buffer, and incubated for 1.5 hours at 4 °C on a turning wheel (Fisherbrand™ Multi-Purpose Tube Rotators). The samples were loaded on Micro Bio-Spin Columns columns (Bio-Rad) pre-equilibrated in cell opening buffer, and washed 10 times with 1 mL of cell opening buffer by gravity flow. RqcH-GS-FLAG3 was eluted by addition of 200 μL opening buffer containing 0.1 mg/mL poly-FLAG peptide (Biotool, Bimake) for 45 min on a turning wheel. All incubations, washes and elutions were performed at 4 °C. The eluted sample was collected by centrifugation at 2000 rpm for 1 minutes 4 °C in a F241.5P rotor using a 149 Microfuge 22R centrifuge (Beckman Coulter). One aliquot of the eluted sample was resolved on SDS-PAGE, the other was blotted on cryo-EM grids, and the remaining sample was used for mass spectrometry and tRNA-array analyses. For SDS-PAGE analyses, 20 μL aliquots of samples (flowthrough, washes and elutions) were mixed with 5 μL of 5x SDS loading buffer and heated at 95 °C for 15 minutes. The beads remaining in the column were washed twice with 1 mL of cell opening buffer and resuspended in 100 μL of 1x SDS loading buffer. Denatured samples were loaded on 12% SDS-PAGE. SDS-gels were stained by “Blue-Silver” Coomassie Staining (Candiano et al., 2004) and washed with water for 6 hours or overnight before imaging with LAS4000 (GE Healthcare).

### Preparation of cryo-EM grids

Eluted pull-down samples were kept on ice and loaded on grids within two hours after preparation without freezing. The concentration of ribosomes in the samples was estimated from SDS-PAGE gels by comparison of ribosomal band intensities in eluted samples with the bands from loaded ribosomes with known concentration. The concentration of ribosomes in elution of RqcH-FLAG3 and YabO-FLAG3 was about 20 nM and 100 nM, respectively. Vitrobot (FEI) blotting was performed at 100% humidity, 4 °C, 5 seconds blot time, 1 second wait time and 0 second drain time; the resultant sample was vitrified by plunge-freezing in liquid ethane. Grids were imaged on a Titan Krios (FEI) operated at 300 kV at a nominal magnification of 165 000× and a pixel size of 0.82 Å with a Gatan K2 Summit camera with a 4 seconds exposure and 20 frames using the EPU software. For RqcH-FLAG3 pulldowns, two data sets were collected on Quantifoil 2/1 Cu 300 and Quantifoil 2/2 Cu 300 grids. The YabO-FLAG3 data set was collected on a carbon-coated Quantifoil 2/2 Cu 300 grid.

### Cryo-EM data processing

Processing was performed with Relion 3.1 unless otherwise stated (Zivanov et al., 2018). Movies were aligned with MotionCor2 with 5 × 5 patches (Zheng et al., 2017) and the CTF was estimated with Gctf (Zhang, 2016). Particles were picked with crYOLO using the provided general model (Wagner et al., 2019), and initially extracted with a box size of 140 pixels, pixel size of 2.46 Å. 3D classifications were performed without angular sampling. For focused classification with partial signal subtraction, the volume eraser and vop commands in UCSF Chimera were used to create starting volumes for masks. For high-resolution refinements, particles were re-extracted in a box of 420 pixels with a pixel size of 0.82 Å. For CTF refinement, anisotropic magnification, higher order aberrations, and per-particle defocus and astigmatism were refined. Volumes were locally filtered with SPHIRE (Moriya et al., 2017), and local resolution was estimated with ResMap (Kucukelbir et al., 2014). The pixel size of the final maps was estimated by comparison to existing structures using UCSF Chimera. Resolutions were estimated with RELION using the ‘gold standard’ criterion (Scheres and Chen, 2012).

For the RqcH IP sample, 725,554 particles were initially picked from 6,730 micrographs (selected from 7,177 initial micrograph movies) from two separate data collections. After 2D classification, 724,098 particles were selected for further processing. An initial model was made *de novo* using the RELION 3D initial model tool, and this was low-pass filtered to 60 Å and used as a reference for a 3D refinement of all particles selected after 2D classification. 3D classification with eight classes was then performed without alignment. The two classes from this classification that contained 50S, RqcH and tRNA, but no extra density towards the edge of the volume (643,616 particles or 88.9% of the starting particles) were selected for further subsorting. 3D refinement was repeated, a generous soft mask encompassing the A-, P- and E-sites was used for partial signal subtraction, and 3D classification was performed with eight classes, T = 200 and the resolution of the expectation step limited to 10 Å. Two classes (totalling 13.9% of the particles) contained RqcH and an A/P-tRNA, with the class among these that had the most interpretable density for the RqcH NFACT-N and HhH domains was selected for further subsorting. Partial signal subtraction around the A- and P-sites was followed by 3D classification with four classes, T = 200, and the resolution of the expectation step limited to 10 A. The class with the most interpretable density for the RqcH NFACT-N and HhH domains (totalling 10,703 particles or 24.3% of the total particles) was selected for 3D refinement and designated State A. Three classes (totalling 51.5% of the particles) contained RqcH and P-site tRNA. The class with the most interpretable density (containing 74,210 particles) was chosen for further refinement and the resulting volume designated State B. A class with particularly poorly-resolved RqcH NFACT-N and HhH domains (containing 110,597 particles) was designated State B*. The third class resembled state B with the RqcH NFACT-N and HhH domains modestly shifted away from the P-site tRNA ASL. A class with 6.1% of particles (39,077 total) contained an E-site tRNA, and was designated State C. A class containing 28.4% of particles (182,833 total) contained P-site tRNA and YabO, but no RqcH, and was designated State D. The resolution of this volume was enhanced by CTF refinement. The final remaining class, containing 10.3% of particles, contained density corresponding to the RqcH CC-M, but not the NFACT-N or HhH, domains.

For the YabO IP sample, 592,872 particles were initially picked from 5,242 micrographs (selected from 5,614 initial micrograph movies). After 2D classification, 579,606 particles were selected for further processing. State D from the RqcH IP processing was low-pass filtered to 60 Å and used as a reference for 3D refinement prior to 3D classification with eight classes and no angular sampling. Four classes comprising 98.3% of particles (569,758 total) were recognisably 50S and were selected for further processing. 3D refinement was repeated, a generous mask around the A-, P- and E-sites was used for partial signal subtraction, and 3D classification was performed with eight classes, T = 200 and the resolution of the expectation step limited to 10 Å. Four of the resulting classes, comprising 31.1% of starting particles (182,833 total) resembled State D and were refined further, including CTF refinement. One class comprising 9.3% of particles (53,124 total) resembled State B, and another with 8.0% (45,313 particles) and which contained P-site tRNA, E-site tRNA and YabO was designated State E. The remaining two classes consisted of an apparent 50S with no ligand (12.8%) or with poorly resolved RqcH (14.1%) and were not refined further. For RsfS-focussed classification, a soft mask around RsfS was used for partial signal subtraction, and 3D classification was performed with four classes, T = 50 and the resolution of the expectation step limited to 10 A. Starting models for the 60S subunit were taken from PDB entries 6HA1 and 6HA8, as well as 4V9F for the uL11/H44 stalk base (Crowe-McAuliffe et al., 2018; Gabdulkhakov et al., 2013). For RqcH and YabO, SWISS-MODEL (Waterhouse et al., 2018) was used to generate homology models using the following templates: RqcH NFACT-N, 6PON (Manne et al., 2019); RqcH HhH, 3DOA and 6PON; RqcH CC and NFACT-R domains, 5H3W (Musyoki et al., 2016); YabO, 1DM9 (Staker et al., 2000). PDB entries 5H3X and 3J92 were additionally used to help with modeling RqcH (Musyoki et al., 2016; Shao et al., 2015). PDB entry 1EHZ was used as a template for modelling *B. subtilis* alanine tRNA-TGC-1-1 (Shi and Moore, 2000).

Models were initially fitted with UCSF Chimera (Pettersen et al., 2004) or aligned with Pymol (Schrödinger, https://pymol.org), and manually adjusted with Coot (Emsley et al., 2010). The initial RqcH model was built using the volume from multibody refinement, and individual domains from this model were then placed in the other maps with minor adjustments. Serine 2 was chosen as the starting amino acid because a peptide lacking the initiator methionine was the most abundant in mass spectrometry (**Table S1**). The linker regions between the NFACT-N and HhH domains (residues 174–178), as well as between the CC-M and NFACT-R domains (residues 434–445), were poorly resolved and therefore not included in the final model. The NFACT-R domain was particularly poorly resolved and was therefore modelled as poly-alanine only. For the 50S ribosomal subunit, the uL1 stalk and tip of the ASF were flexible and were not included in the final models. The YabO model was built initially into the State B volume. Phenix was used for refinement (Liebschner et al., 2019). States A and B were refined against locally filtered volumes. The RqcH-focused State B multibody refinement was refined against a volume that had been sharpened using the RELION post-processing procedure with automatic b-factor estimation.

### tRNA microarrays

tRNA microarrays were performed similarly to as previously described (Beckert et al., 2018). The RqcH-50S-bound tRNA (i.e. in the immunoprecipitated RqcH-FLAG_3_ aliquots) was compared on the same arrays to the total *B. subtilis* tRNA. A detailed protocol is published on protocols.io (dx.doi.org/10.17504/protocols.io.hfcb3iw). For deacylation lysate and immunoprecipitated samples were incubated with 125 mM Tris-HCl, pH = 9.0, 0.1 M EDTA, 0.5% (w/v) SDS at room temperature for 45 minutes, before neutralisation with an equal volume of 1 M NaOAc, pH = 5.5. RNA was extracted twice with 5:1 acidic phenol:chloroform, precipitated with ethanol, and resuspended in ddH_2_O. Using the unique invariant single stranded 3’-NCCA-ends of intact tRNA a Cy3-labeled RNA/DNA and CAtto647-labeled RNA/DNA hybrid oligonucleotide was ligated to the tRNA extracted from the RqcH-50S samples and total *B. subtilis* tRNA, respectively. Labeled RNA was purified by phenol:chloroform extraction and ligation efficiency verified on denaturing 10% SDS-PAGE. Labeled tRNA samples were loaded on a microarray containing 24 replicates of full-length tDNA probes recognizing 36 *B. subtilis* tRNA isoacceptors and hybridized for 16 h at 60°C. Fluorescence signals of microarrays were recorded with a GenePix 4200A scanner (Molecular Devices) and statistically analyzed with in-house scripts with Python version 3.7.0. Data have been deposited in Gene Expression Omnibus (GEO) database under accession GSE152592.

### Proteomics sample preparation and LC/MS/MS analysis

Proteins were precipitated with 10% (w/v) trichloroacetic acid overnight at 4 °C, pelleted at 17,000 g 4 °C and washed twice with cold 90% (v/v) acetone. Precipitated proteins were solubilized in 7 M urea, 2 M thiourea, 100 mM ammonium bicarbonate (ABC) buffer, reduced with 5 mM dithiothreitol for 30 min at room temperature (RT) and alkylated with 20 mM chloroacetamide in the dark. Pre-digestion with 1:50 (enzyme to protein ratio) *Lysobacter enzymogenes* Lys-C (Fujifilm Wako Pure Chemical) was carried out for 4 hours at RT. Next, the solution was diluted five times with 100 mM ABC buffer and a further digestion with 1:50 dimethylated *Sus scrofa* trypsin (Sigma Aldrich) was carried out overnight at RT. Samples were then acidified with trifluoroacetic acid (TFA) added to 1.0% (v/v), and desalted on in-house made C18 SPE tips. Purified peptides were reconstituted in 0.5% TFA (v/v) for nano-LC/MS/MS.

Peptides were injected to an Ultimate 3000 RSLCnano system (Dionex) using a 0.3 × 5 mm trap-column (5 μm C18 particles, Dionex) and an in-house packed (3 μm C18 particles, Dr Maisch) analytical 50 cm × 75 μm emitter-column (New Objective). Peptides were eluted at 250 nL/min with an 8-40% (2 h) A to B gradient (buffer A: 0.1% (v/v) formic acid; buffer B: 80% (v/v) acetonitrile + 0.1% (v/v) formic acid) to a quadrupole-orbitrap Q Exactive Plus (Thermo Fisher Scientific) MS/MS via a nano-electrospray source (positive mode, spray voltage of 2.5 kV). The MS was operated with a top-5 data-dependent acquisition strategy. Briefly, one 350-1,400 m/z MS scan at a resolution setting of R = 70,000 was followed by higher-energy collisional dissociation fragmentation (normalized collision energy of 26) of the 5 most intense ions (z: +2 to +6) at R = 17,500. MS and MS/MS ion target values were 3,000,000 and 50,000 ions with 50 and 100 ms injection times, respectively. Dynamic exclusion was limited to 40 s.

MS raw files were processed with the MaxQuant software package (version 1.6.1.0) (Tyanova et al., 2016). Methionine oxidation, protein N-terminal acetylation, protein N-terminal methionine formylation and removal of up to 4 N-terminal amino acids were set as potential variable modifications, while cysteine carbamidomethylation was defined as a fixed modification. Identification was performed against the UniProt (www.uniprot.org) database (*B. subtilis* wild-type strain 168, 4 271 protein sequences) using the tryptic digestion rule (i.e. cleavages after lysine and arginine without proline restriction). Only identifications with at least 1 peptide ≥ 7 amino acids long (with up to 2 missed cleavages) were accepted. Label-free intensity normalization with the MaxLFQ algorithm (Cox et al., 2014) was also applied. Protein and LFQ ratio count (*i.e*. number of quantified peptides for reporting a protein intensity) was set to 1. iBAQ feature of MaxQuant was enabled. This normalizes protein intensities by the number of theoretically observable peptides and enables rough intra-sample estimation of protein abundance. Peptide-spectrum match, peptide and protein false discovery rate was kept below 1% using a target-decoy approach (Elias and Gygi, 2007). All other parameters were default.

The mass spectrometry raw files along with MaxQuant identification and quantification outputs (txt folder) have been deposited to the ProteomeXchange Consortium (Vizcaino et al., 2014) via the PRIDE partner repository with the dataset identifier PXD019364.

### Alignment and phylogenetic analysis

For the sequence alignment and secondary structure assignment in **Figure 3D**, YabO from State B and Hsp15 from PDB 1DM9 were aligned by structure with the DALI server (Holm, 2019) and DSSP was used to annotate secondary structural elements. For the C-terminal extension of Hsp15, no structural information is available and PSI-PRED was used to predict secondary structure (Buchan and Jones, 2019).

YabO and RqcH sequences were retrieved from the NCBI protein database, using accession numbers from the COG 2014 database (Galperin et al., 2015). The YabO/Hsp15/RluA group belongs to COG1188 (411 sequences), and RqcH to COG1293 (357 sequences). Sequences were aligned using MAFFT-L-INS-I v6.861b (Katoh et al., 2005), including curation to remove 55 non-alignable sequences in the case of COG1293 and 29 RluA sequences (more distant relatives of YabO/Hsp15 but also carrying the S4 domain) in the case of COG1188. Representative YabO/Hsp15 sequences were selected for phylogenetic analysis for **Figure 3D** to sample broadly across protein diversity and taxonomic distributions. After trimming the alignment to remove columns with <50% gaps with TrimAL v1.4 (Capella-Gutierrez et al., 2009), phylogenetic analysis was carried out with RaxML v 8.2.12 (Stamatakis, 2014) on the Cipres Science Gateway (Miller et al., 2015) with 100 bootstrap replicates and the LG model of substitution.

### Figure preparation

Figures were prepared using UCSF ChimeraX (Goddard et al., 2018) and Inkscape (https://inkscape.org/).

## AUTHOR CONTRIBUTIONS

D.N.W. and V.H. designed the study. H.T. and V.M. prepared the cryo-EM samples. H.T. performed biochemical and genetic studies. C.C.-M. processed the cryo-EM data, built and refined the molecular models. S.K. and T.T. performed the mass spectrometry analysis. G.C.A. performed sequence and phylogenetic analysis. C.P and Z.I. performed the tRNA microarray analysis. All authors interpreted the results and helped D.N.W. and C.C.-M. write the paper.

## ACKNOWLEDGMENTS

We thank Michael Hall for help with cryo-EM data collection. The electron microscopy data was collected at the Umeå Core Facility for Electron Microscopy, a node of the Cryo-EM Swedish National Facility, funded by the Knut and Alice Wallenberg, Family Erling Persson and Kempe Foundations, SciLifeLab, Stockholm University and Umeå University. This research was supported by grants from the Deutsche Forschungsgemeinschaft WI3285/8-1, SPP-1879 (to D.N.W.), the Swedish Research Council (Vetenskapsrådet) grants (2017-03783 to V.H. and 2019-01085 to G.C.A.), Ragnar Söderbergs Stiftelse (to V.H.), postdoctoral grant from the Umeå Centre for Microbial Research, UCMR (to H.T.), the European Union from the European Regional Development Fund through the Centre of Excellence in Molecular Cell Engineering (2014-2020.4.01.15-0013 to T.T. and V.H.); and the Estonian Research Council (PRG335 to T.T. and V.H.). D.N.W. and V.H. groups are also supported by the Deutsche Zentrum für Luft- und Raumfahrt (DLR01Kl1820 to D.N.W.) and the Swedish Research Council (2018-00956 to V.H.) within the RIBOTARGET consortium under the framework of JPIAMR.

## SUPPLEMENTAL INFORMATION

### SUPPLEMENTAL FIGURES

**Figure S1.**
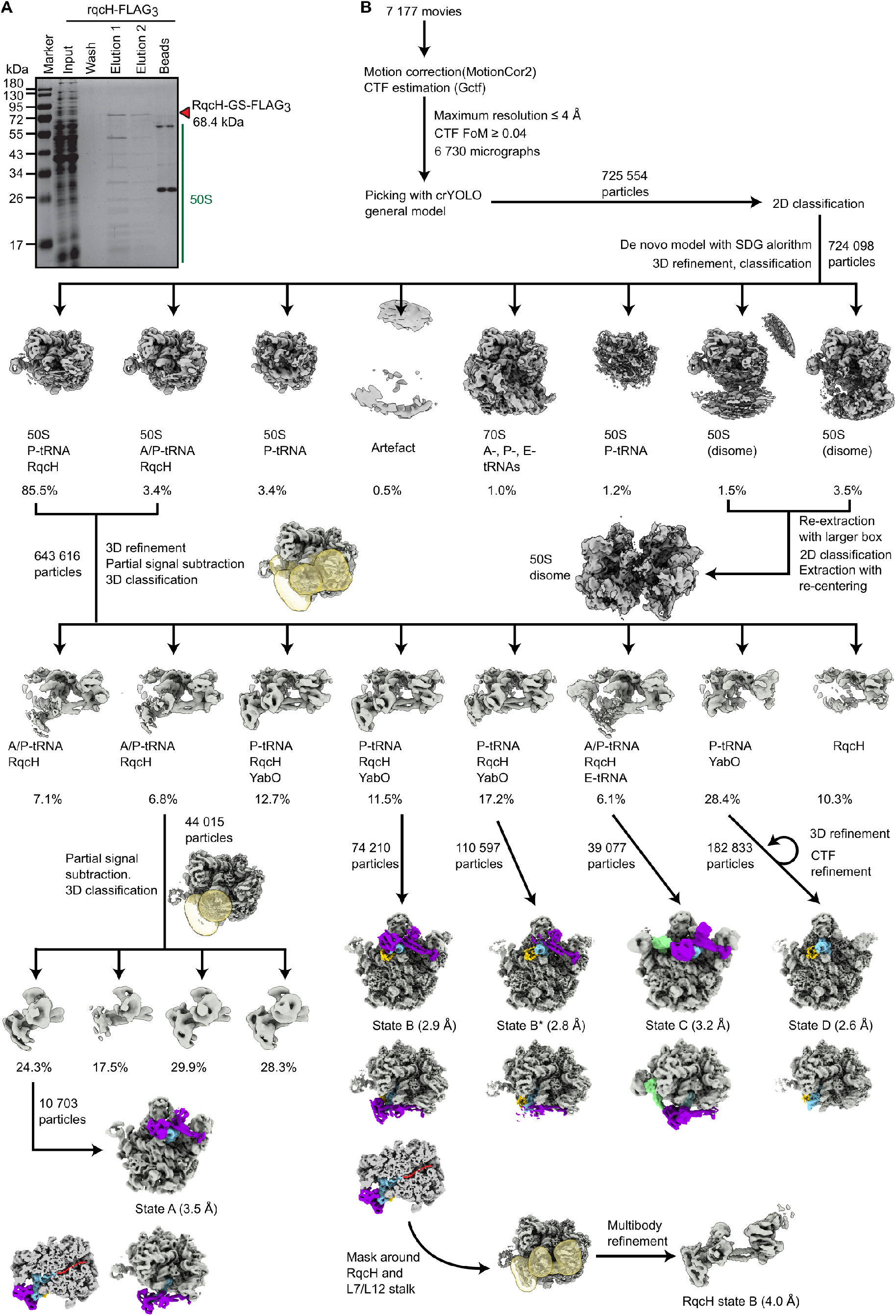
Processing of cryo-EM data from RqcH–FLAG immunoprecipitation. (A) Immunoprecipitation of RqcH-FLAG3. (B) Processing of RqcH IP micrographs. Refer to methods for additional details.

**Figure S2.**
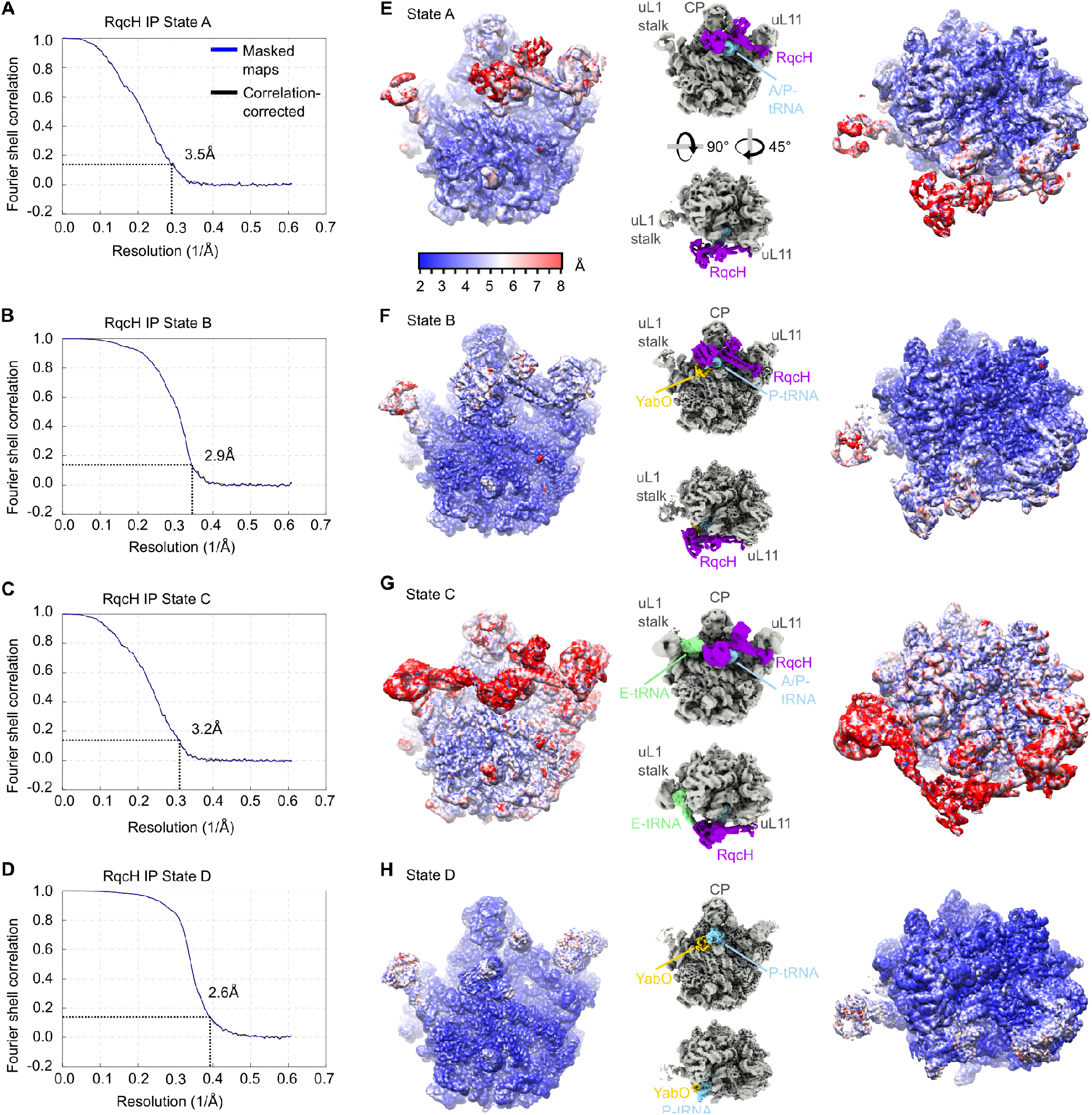
Average and local resolution of cryo-EM maps from RqcH–FLAG immunoprecipitation. (A-D) FSC curves generated by RELION for each class. The dashed line indicates an FSC of 0.143. (E-H) EM maps of each state colored according to local resolution, with small insets colored as in Fig. 1A-C to illustrate the orientation of the particles.

**Figure S3.**
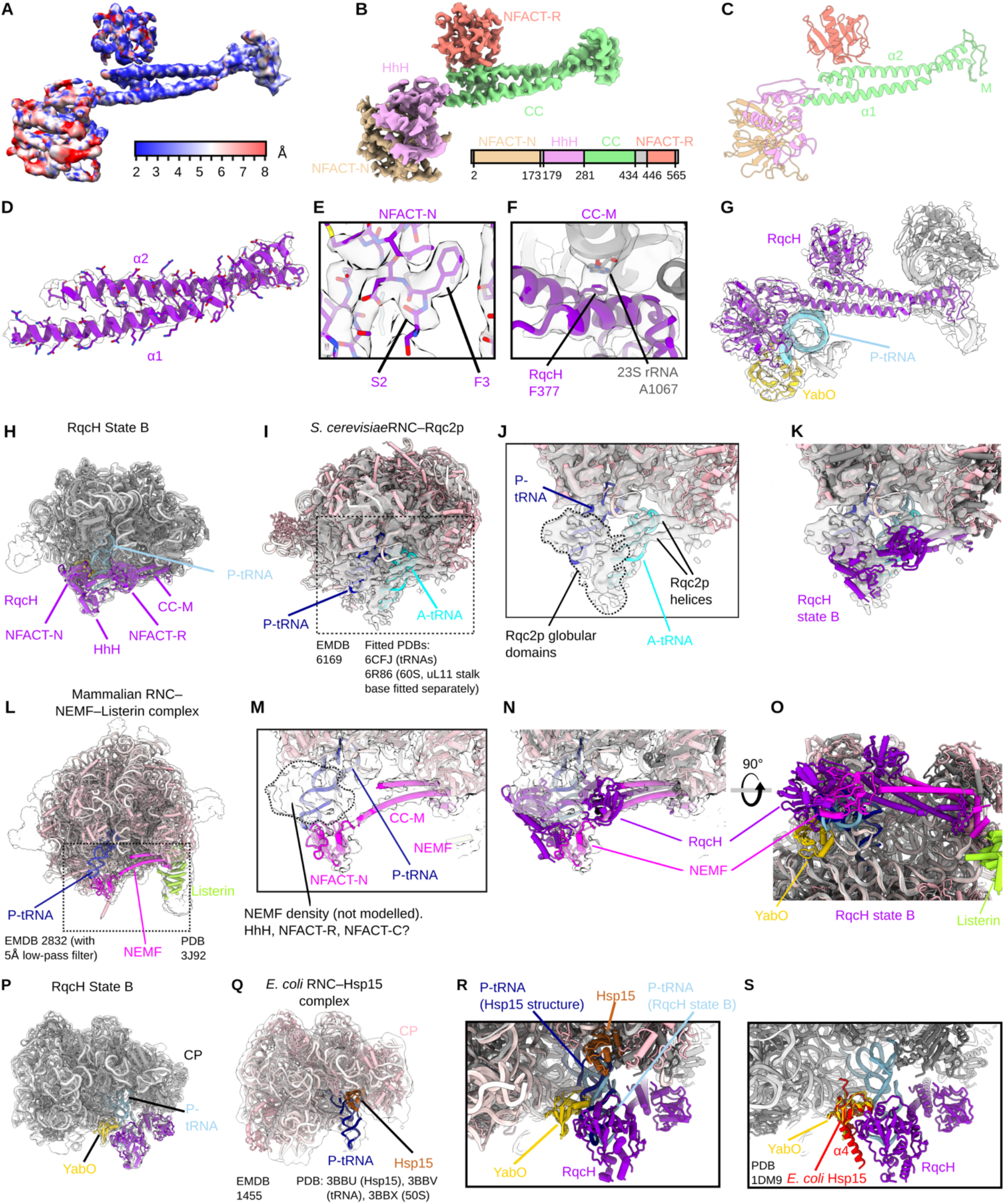
State B RqcH-focused multibody refinement and comparison with eukaryotic Rqc2/NEMF-ribosome structures. (A-C) RqcH from State B multibody refinement with cryo-EM map colored according to (A) local resolution, (B) domain, or (C) model only. (D) helices of the RqcH CC-M domain. (E) The RqcH N-terminus. (F) View of RqcH F377, located in α2, interacting with the 23S rRNA. (G) Components of the multibody refined map (transparent grey) with fitted model. (H-I) Overview of RqcH State B and the *S. cerevisiae* Rqc2p–RNC complex. To aid interpretation, models of the yeast 60S subunit (PDB 6R86) and A- and P-tRNAs (from PDB 6CFJ) were fitted into density from EMD-6169 (Shen et al., 2015; Su et al., 2019; Tereshchenkov et al., 2018). The uL11/H44 stalk base from PDB 6R86 was fitted independently from the rest of the 60S. No model is available for Rqc2p. (J-K) Close views of (J) Rqc2p with positions of the Rqc2p globular domains (dotted lines) and helices indicated, with (K) RqcH State B overlaid. (L) Overview of mammalian RNC–NEMF–Listerin complex with partial NEMF model (Shao et al., 2015). (M-O) Close view of NEMF, with density corresponding to unmodelled NEMF globular domains indicated, with (N) RqcH from State B overlaid, and (O) a rotated view with models only to compare conformations of NEMF and RqcH, coloured as in (H) and (I). (P-Q) Views of RqcH State B (P) and *E. coli* RNC–Hsp15 (Q) complexes. (R) Overlay of the two models showing YabO and Hsp15 binding. (S) Hsp15 from *E. coli* (PDB 1DM9) (Staker et al., 2000) aligned with YabO from RqcH State B. C-terminal helix α4, which is not present in YabO, is indicated.

**Figure S4.**
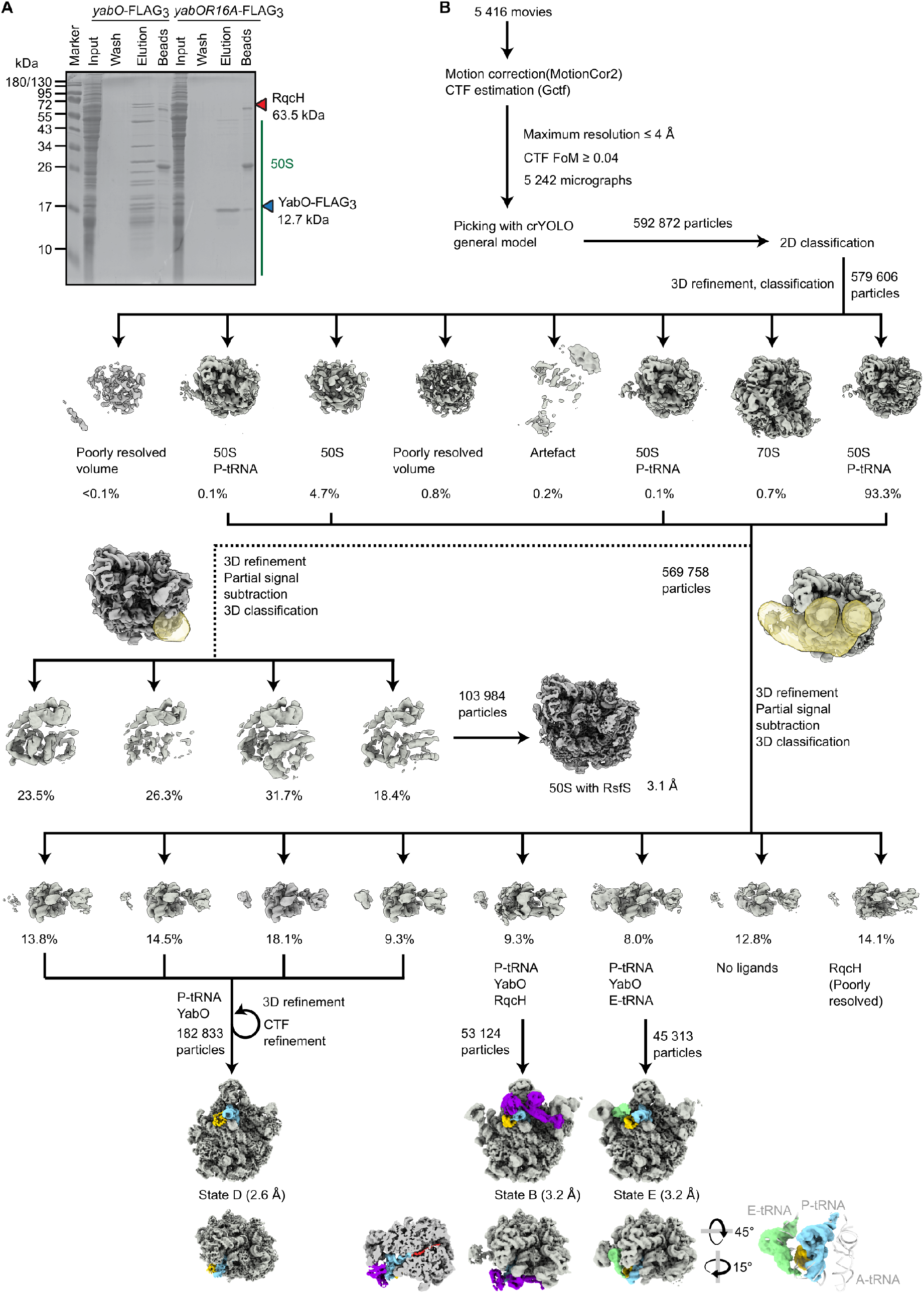
Processing of cryo-EM data from YabO–FLAG3 immunoprecipitation. (A) Immunoprecipitation of C-terminal FLAG-tagged YabO or YabO R16A mutant. (B) Processing of cryo-EM data from YabO–FLAG3 immunoprecipitation. Refer to methods for additional details.

**Figure S5.**
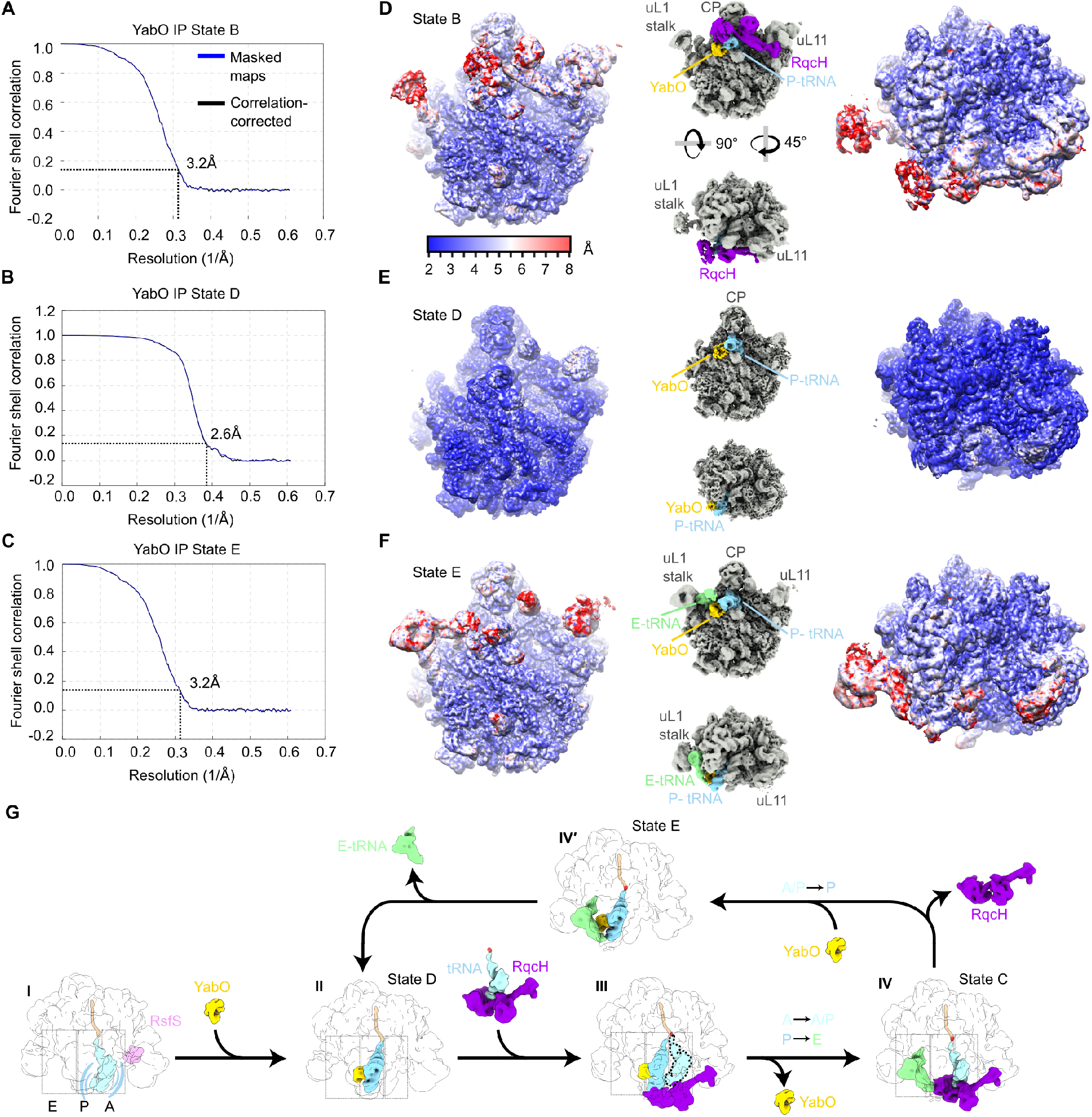
Average and local resolution of cryo-EM maps from YabO–FLAG immunoprecipitation. (A-C) FSC curves generated by RELION for YabO pull-out (A) State B, (B) State D and (C) State E. The dashed line indicates an FSC of 0.143. (D-F) EM maps of each state colored according to local resolution, with insets colored as in Fig. 1A-C to illustrate the orientation. (G) Alternative branch of scheme in Figure 6 with IV’, corresponding to State E, observed only in the YabO immunoprecipitation.

**Figure S6.**
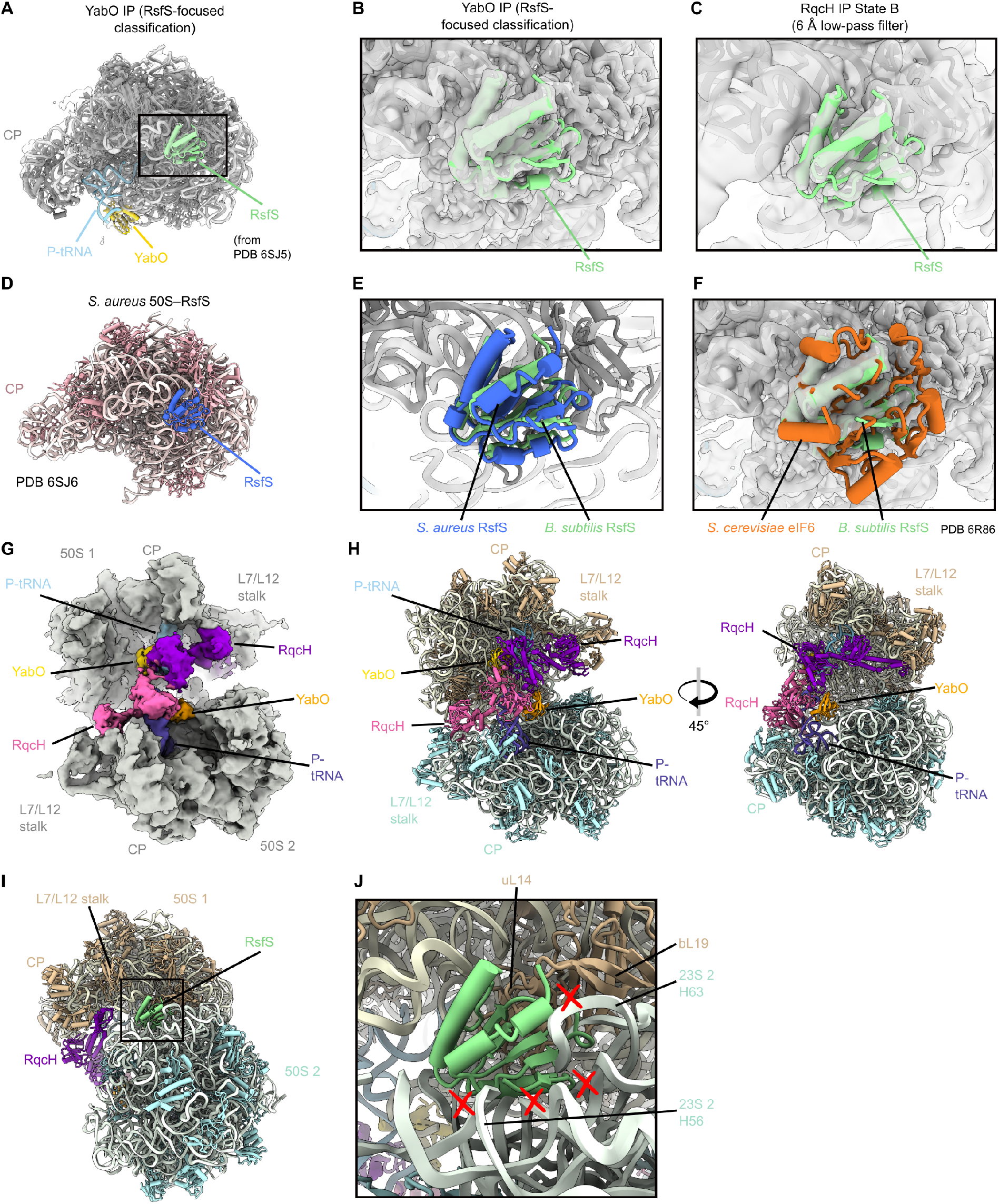
RsfS binds 50S to both RqcH- and YabO-associated 50S. (A-B) YabO RsfS-focused structure with fitted RsfS from PDB 6SJ5 (Khusainov et al., 2020). (C) Same view as in d except with a low-pass filtered RqcH State B map. (D) RsfS bound to the *S. aureus* ribosome. (E-F) Overlay of *S. aureus* RsfS (blue, E) or eIF6 (F) (Su et al., 2019) on the RsfS-bound *B. subtilis* 50S. (G-H) The 50S disome observed in the RqcH pulldown, resembling two State B particles bound via the intersubunit interface. (I-J) RsfS, as in panel (C), superimposed on the 50S disomes.

